# The Kendrick Modelling Platform: Language Abstractions and Tools for Epidemiology

**DOI:** 10.1101/289199

**Authors:** Mai Anh T. Bui, Nick Papoulias, Serge Stinckwich, Mikal Ziane, Benjamin Roche

## Abstract

**Background:** Mathematical and computational models are widely used for examining transmission, pathogenicity and propagation of infectious diseases. Software implementation of such models and their accompanied modelling tools do not adhere to well established software engineering principles. These principles include both *modularity* and *clear separation of concerns* that can promote *reproducibility* and *reusability* by other researchers. On the contrary the software written for epidemiology is monolithic, highly coupled and severely heterogeneous. This reality ultimately makes these computational models hard to study and to reuse, both because of the programming competence required and because of the incompatibility of the different approaches involved. Our goal with **Kendrick** is to simplify the creation of epidemiological models through a unified Domain-Specific Language for epidemiology that can support a variety of modelling and simulation approaches classically used in the field. This goal can be achieved by promoting *reproducibility* and *reuse* with modular modelling abstractions.

**Results:** We show through several examples how our modular abstractions and tools can reproduce uniformly complex mathematical and computational models of epidemics, despite being simulated by different methods. This is achieved without requiring sophisticated programming skills from the part of the user. We then successfully validate each kind of simulation through statistical analysis between the time series generated and the known theoretical expectations.

**Conclusions:** **Kendrick** is one of the few DSLs for epidemiology that does not burden its users with implementation details or expecting sophisticated programming skills. It is also currently the only language for epidemiology that supports modularity through clear separation of concerns that promote reproducibility and reuse. **Kendrick**’s wider adoption and further development from the epidemiological community could boost research productivity in epidemiology by allowing researchers to easily reproduce and reuse each other’s software models and simulations. The tool can also be used by people who are not necessarily epidemiology modelers.

## Background

During the last decades, mathematical and computational modelling became widely used for investigating the mechanisms of infectious diseases propagation [1, 2], exploring their evolutionary dynamics [3, 4], and/or informing control strategies [5, 6]. Their insights are increasingly recognised since the early 1980s when human population entered a period of emergence and re-emergence of infectious diseases [7, 8]. Including the increasing number of public health threats, such as a possible small-pox epidemic due to bioterrorism attack [9], emergence of pandemic influenza [10] or the need to propose new kinds of vaccination programs [11]. Indeed, the complexity of these new infections, relying on many inter-connected factors [12, 13, 14] such as biodiversity decline [15], antibiotics resistance [7] or intensification of worldwide trade [16], makes the anticipation of their propagation and their evolution a very tricky challenge against which epidemiological models can be crucial tools.

Epidemiological models largely rely on the so-called SIR framework [17, 18] where host population is divided into categories corresponding to their epidemiological status. Basically, epidemiological models consider individuals who are Susceptible to the pathogen (state *S*) and then can be infected, Infectious (state *I*) that can transmit the disease and Recovered (state *R*) who are immunised and cannot become infected again. Such model aims to characterise the transition between these categories and the consequences on the dynamics of each category, generally the ‘Infectious’ one that contains diseased individuals. From this initial configuration, an infinity of other categories can be added in order to represent different transmission cycles corresponding to each infectious disease that could be studied.

Based on this framework, an epidemiological model can nevertheless be implemented and simulated in different ways. Generally, the first approach is to define the life cycle through a system of ordinary differential equations in order to be analytically tractable or to be deterministically simulated through numerical methods, such as Runge-Kutta [19]. While this approach is especially useful to understand the average dynamics without chance, shifting to a stochastic approach (e.g., through Gillespie algorithms [20]) is known to be more realistic [18] and can significantly impact the simulated dynamics of infectious diseases (such as their seasonality [18]). Finally, an agent-based implementation is sometimes required to reach a level of detail that would not be tractable with other approaches because of combinatorial explosion [21].

One of the problems of modelling is bridging the gap between conceptual models (categories and their transition) and their computer simulation (through deterministic, stochastic or agent-based implementation). In fact, going from a conceptual model to a computational one requires some programming skills, which could be very rudimentary for the deterministic implementation to extremely complex for agent-based models. However, this gradient of programming complexity yields a high heterogeneity in the code that could be produced, with non negligible impacts on the disease dynamics. For instance, despite their simplicity, deterministic models can show very different dynamics according to the algorithm being used [18]. We also observe that the software implementation of such models and their accompanied modelling tools do not adhere to well established software engineering best practices such as *modularity* through *clear separation of concerns* that can promote *reproducibility* and *reusability* by other researchers.

Domain Specific Languages (DSLs) address such difficulties by separating modelling and specification concerns (conceptual model) from implementation aspects (computational model). Contrary to General-purpose Programming Languages (GPLs), DSLs are higher-level languages that provide a more expressive syntax based on abstractions and notations that are directly-related to the concepts of the studied domain [22][23][24].

Our goal with **Kendrick** is to simplify the creation of epidemiological models through a unified Domain-Specific Language for epidemiology that can support a variety of modelling and simulation approaches used in the field. This goal can be achieved by promoting *reproducibility* and *reuse* with modular modelling abstractions.

**Kendrick** is designed according to a general meta-model that captures the basic domain concepts and opens up the possibility of taking into account different aspects of epidemiological modelling. To illustrate the practical usage of our modelling language, we present in this paper two case studies on measles and mosquito-borne diseases. Our approach is carefully validated through statistical comparisons between our simulation results and well-established platform simulations in order to guarantee the link with the mathematical epidemiology literature. While part of the mathematical and syntactical details of the **Kendrick model** and language have been presented before [25][26], this is the first time the software itself is presented, through a step by step coverage of its implementation and usage.

## Implementation

**Kendrick** is implemented on top of the Pharo [27] programming environment using the Moose meta-modelling platform [28]. The numerical analysis back-end is based on the PolyMath framework [29], the visualisation sub-system on Roassal [30, 31] and the UI components on GTTools and the Moldable Inspector [32]. The DSL makes extensive use of Pharo’s reflective sub-system [33], with on-going extensions based on the PetitParser [34] framework and Helvetia parsing workbench [35].

**Kendrick** supports all major desktop platforms (Gnu/Linux, Mac OS and MS Windows) and adopts an open continuous integration process (based on github^[1]^, smalltalkhub^[2]^ and Travis CI^[3]^). It is extensively tested using the SUnit [36] framework, readily available through a dedicated web-site ^[4]^ and well documented through a collaborative wiki ^[5]^.

This section focuses on the implementation details of the **Kendrick** meta-model and DSL. For step-by-step installation and experimentation details see the following Section (*i.e*. Results & Discussion).

### The Kendrick Meta-model

As we saw, **Kendrick** aims to support a wide range of epidemiological models that may use either different formalisms or different implementation techniques.

**Kendrick** needs to abstract away from programming concerns and allow the definition of epidemiological models with little or no programming skills.

In order to achieve these goals we had to identify the proper abstractions and implement them as a general core meta-model, which means implementing: *a set of domain concepts on which all possible base-level models and simulation tools could rely*. The resulting meta-model that we defined turned out to be general enough to capture compartment-based population dynamics rather than just epidemiology.

Seen from this perspective (*i.e*. of compartment-based population dynamics), models aim at answering the following question: *What is the cardinality of the defined compartments at any given time? Or conversely: Given a set of observations, what parameters of an input model can best fit the observed behaviour?*. To answer these questions the **Kendrick** meta-model (seen in Figure 1) includes the following concepts: *Population, Attribute, Equivalence Relation, Compartment, Parameter, Transition and Time Series*.

**Figure 1.**
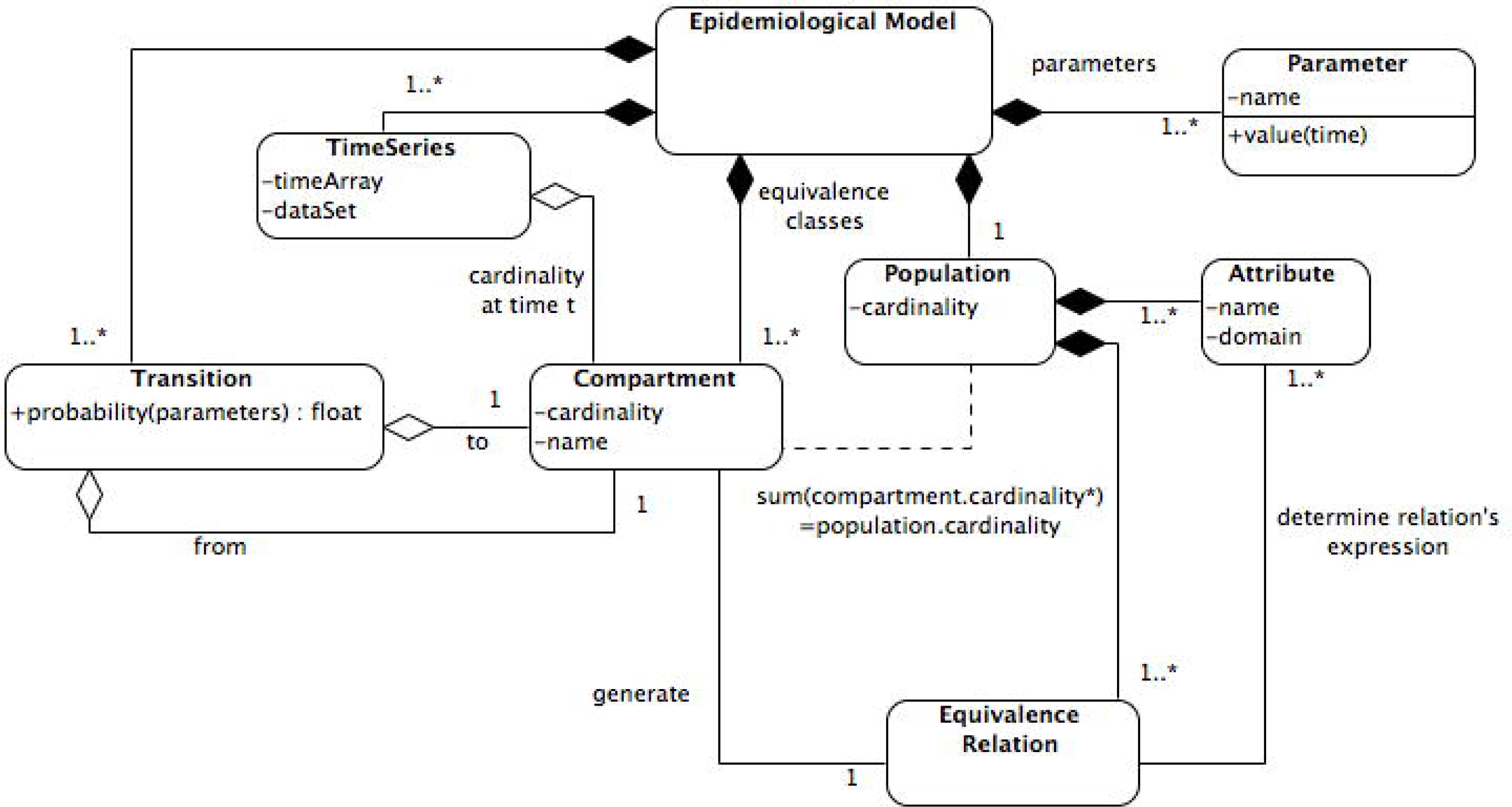
The meta-model UML diagram. This UML diagram shows main elements of our core systems and the relationships among them. Each element in this diagram represents a concept of our language. The composition link between two concepts indicates that the construction of a concept (where the filled diamond of the link points) may involve other ones (at other side of this link).

In order to be as general as possible, *compartments* are assumed to be defined by an *equivalence relationships* on the *population*. In the current version of the meta-model, this is done using the *attributes* of the individuals (such as the infectious status, species, age *etc*.). Moreover, in the implementation, the equivalent individuals are those with the same values for a given set of attributes.

The *Time series* concept captures the simulation results of a model over a certain period of time. Alternatively, partial times series can be used to capture observations. Finally, the defined models may also include additional *parameters* such as: rates, probabilities or initial cardinalities of the population or of compartments. These parameters are captured as numerical temporal functions.

It should be noted here that the **Kendrick** meta-model can capture both deterministic models expressed as a set of ODEs and demographic-stochastic models. Both kinds of models are stored as the transition rate matrix of a continuous-time Markov chain. Different formalisms can be used to write these *transitions*: such as ODEs or stochastic automata (with each syntax being able to be translated into the other). Finally, transition rates are depicted as temporal functions in our meta-model, rather than numbers, to capture variations of these rates (e.g. for seasonal forcing). Any given model can be run as a deterministic, stochastic or agent-based simulation. The default deterministic solver is RK4 [19]. Gillespie’s direct and tau-leap methods are both available for stochastic simulations. Stochastic agent-based simulations can also be run by triggering events at the level of individuals [21].

### The Kendrick DSL

As we saw, the **Kendrick** meta-model tries to address reusability in epidemiological modelling by decomposing highly-coupled monolithic epidemiological models into modular concerns. Indeed, separating these concerns is expected to make them easier to define, understand and change. It is achieved at a mathematical level by defining each concern as a stochastic automaton and by combining them using a tensor sum operator [37]. It is however still necessary to shield epidemiologist from implementation details and avoid to introduce dependencies among concerns in the implementation of models. The goal of the DSL is thus threefold: a) provide an easy-to-use, concise and readable syntax b) hide implementation details as much as possible c) provide a way to keep the concerns separated (i.e. avoid linguistic dependencies between them).

With these goals in mind, our DSL defines the following linguistic (syntactical and semantical) entities: Epidemiological *Models*, *Compositions*, *Scenarios*, *Simulations* and *Visualizations*, which are exemplified in Table 1. As we will see in greater detail in the following section, each one of these entities can be defined as a separate component (in its own file if necessary) and reused several times in multiple projects by simply referring to its name. Moreover, all of the aforementioned entities can be extended (*i.e*. specialized) upon reuse, by simply overriding one or more of their properties. *Models* are the more versatile and reusable entities, since as we can see in the Table below, they define general epidemiology concerns (such as the SIR dynamics or other general aspects, like the type and number of species being modeled).

**Table 1.**
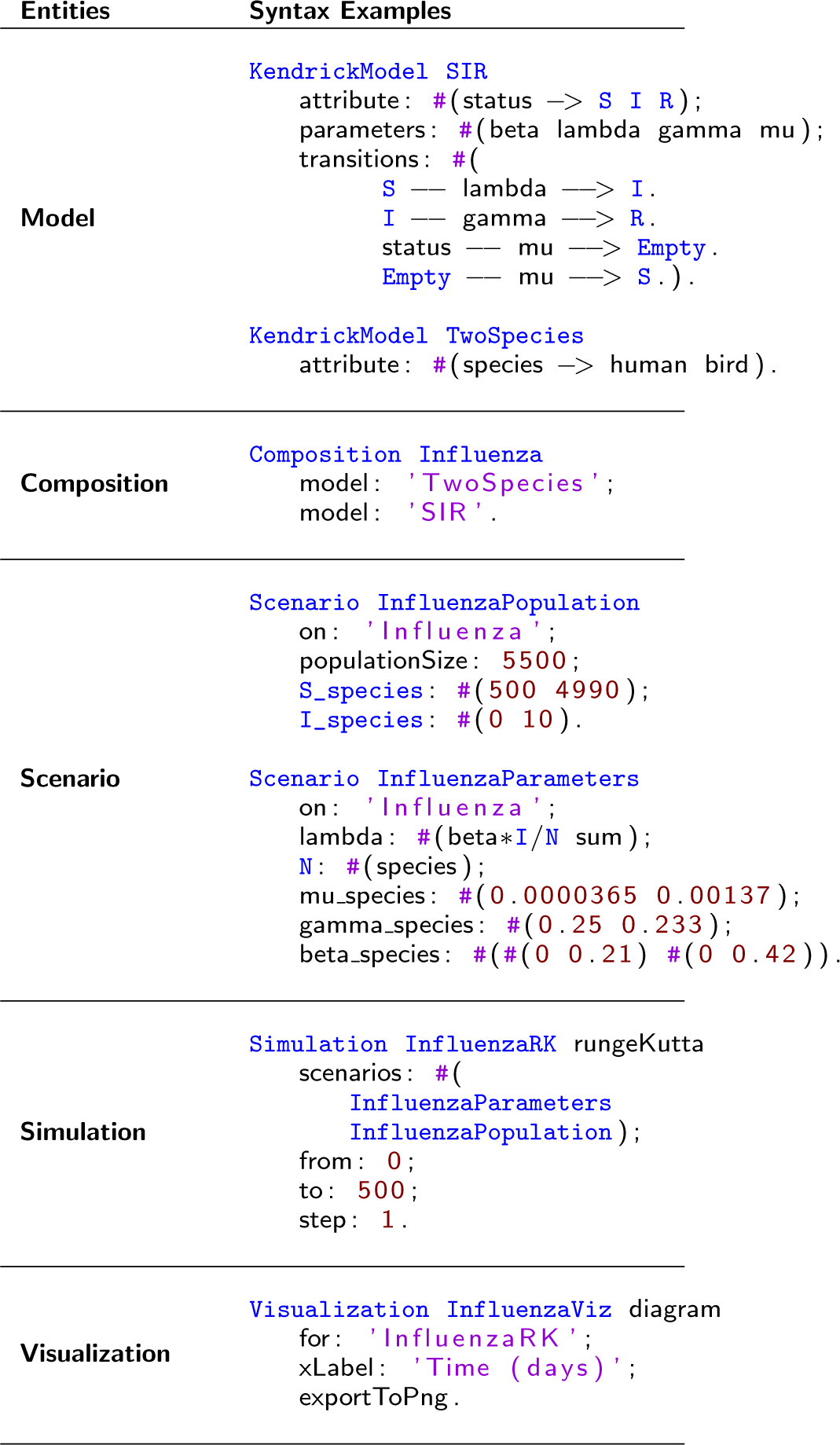
Kendrick DSL Entities (Two species Influenza SIR Example)

These independent models can then be combined (through the *Composition* entity) by further refining cross-concern characteristics. *Scenarios* describe the initialization data of experiments, but can themselves be combined in different ways to provide input to *Simulation* entities. These latter entities define algorithmic properties for experiments, such as solvers and timing constraints. Finally, *Visualization* entities define the desired visual outputs (such as epidemiological figures and maps) that can summarize the experimental results.

## Results & Discussion

In this section, we first provide practical information on how to install and start using **Kendrick**. Then we illustrate in detail how to simulate the evolution of two infectious diseases. The first one is measles, a childhood disease that has been extensively studied through epidemiological modelling. The second one is a vector-borne disease on which we test the multi-host component of our platform.

### Installing & running Kendrick

The easiest way to install **Kendrick** is to download a pre-compiled bundle of the **0.42 version**, for your platform of choice (Mac, Linux and Windows are supported) by following the corresponding links in the Github repository ^[6]^. After unzipping the corresponding .zip file for your platform, you can go ahead and double-click on one of the kendrick launchers (**KendrickUI** for Mac and Linux, or **Kendrick-WinLauncher** for Windows).

Alternatively, we provide a very straight-forward method to compile **Kendrick** from sources on all systems with a bash command-line (this includes Linux, Mac and Windows with Cygwin and/or the Windows 10 Bash sub-system), by issuing the following command:

~~~
wget −O− https://goo.gl/WUQxmp | bash
~~~

This command will automatically retrieve all the required dependencies, compile **Kendrick** from sources and set up all execution scripts for normal or development use. After compiling or downloading the pre-compiled versions of **Kendrick**, the dedicated editor can be run by invoking:

~~~
./KendrickUI
~~~

In which case you will be greeted with the splash screen (Figure 2).

**Figure 2.**
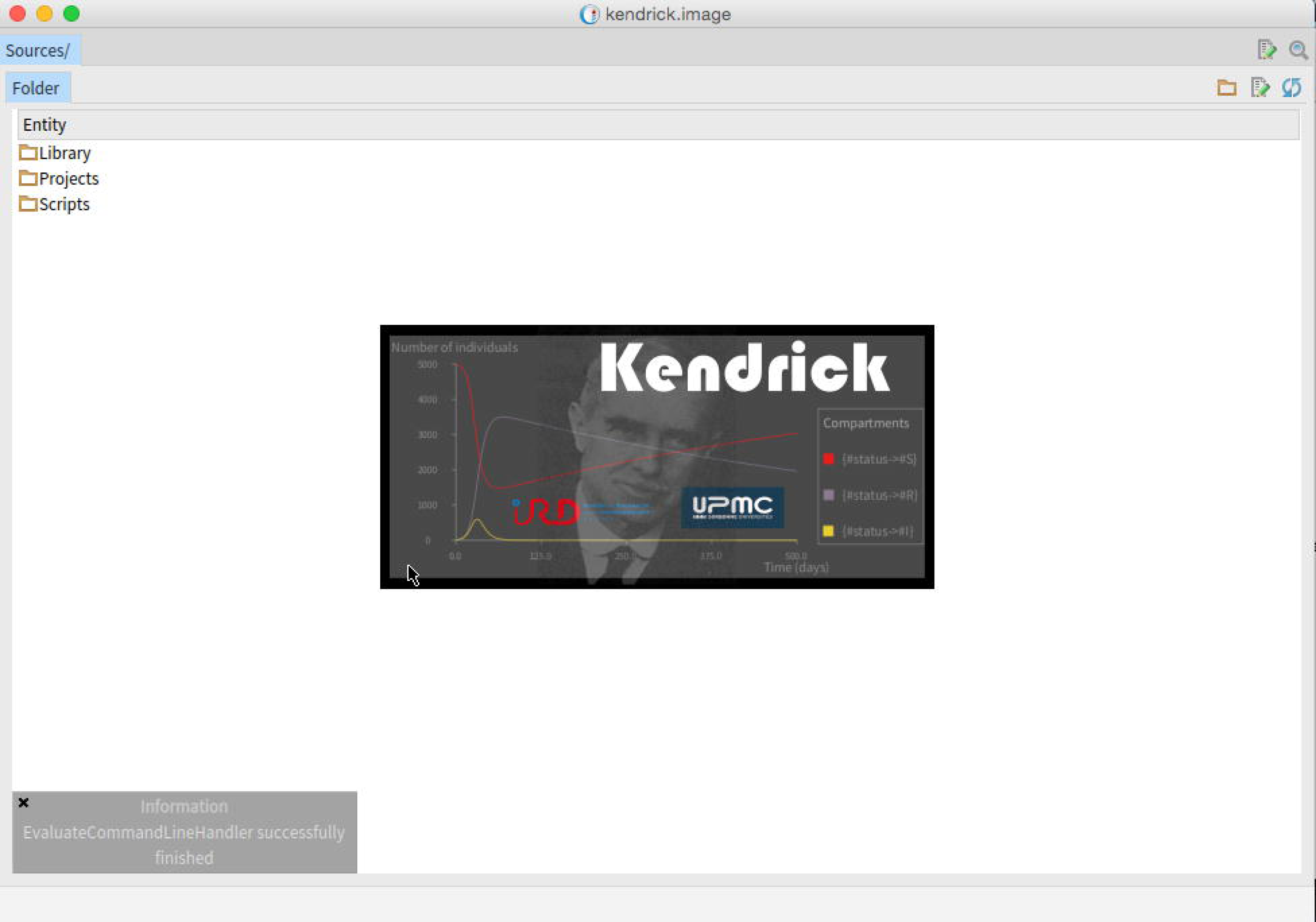
Kendrick UI. Running the Kendrick DSL Editor

Our dedicated editor is a moldable editor / inspector [32] that follows the Cascading Lists ^[7]^ pattern of navigation. This means that each selection reveals a brand new column of actionable information (on the right), with a horizontal bar on the bottom controlling the viewport position and size.

Alternatively, to run **Kendrick** with a full development environment (allowing to use both the DSL and the Pharo API of **Kendrick**), you can invoke:

~~~
./KendrickDevUI
~~~

Finally, to use **Kendrick** with an editor of your choice, you only need to navigate in the Sources directory of your installation, edit or add files for your project and invoke the non-interactive **Kendrick** executable as follows (example for simulating and visualizing the results described in Influenza1Viz.kendrick on your sources folder):

~~~
./Kendrick Sources/Projects/Influenza/Visualization/Influenza1Viz.kendrick
~~~

In this case final visualization results will be produced in:

~~~
Sources/Projects/Influenza/Output/Influenza1Viz.png
~~~

### Case-study I: Measles

We consider the transmission of measles through a SEIR model with demography [38]. Individuals are born with the *Susceptible (S)* status with a birth rate *µ*, then may become *Exposed (E)* (i.e. infected but not yet infectious) with transmission rate *βI*. After an average latent period, 1*/σ*, they may become *Infectious (I)* and may finally switch to *Recovered (R)* after an infectious period: 1*/γ*.

The model is available through the standard **Kendrick** distribution by navigating through our editor either to Scripts/Measles.kendrick (single file script, seen in Figure 3) or to Projects/Measles (several decomposed entities in separate files that are more easily extended and reused).

**Figure 3.**
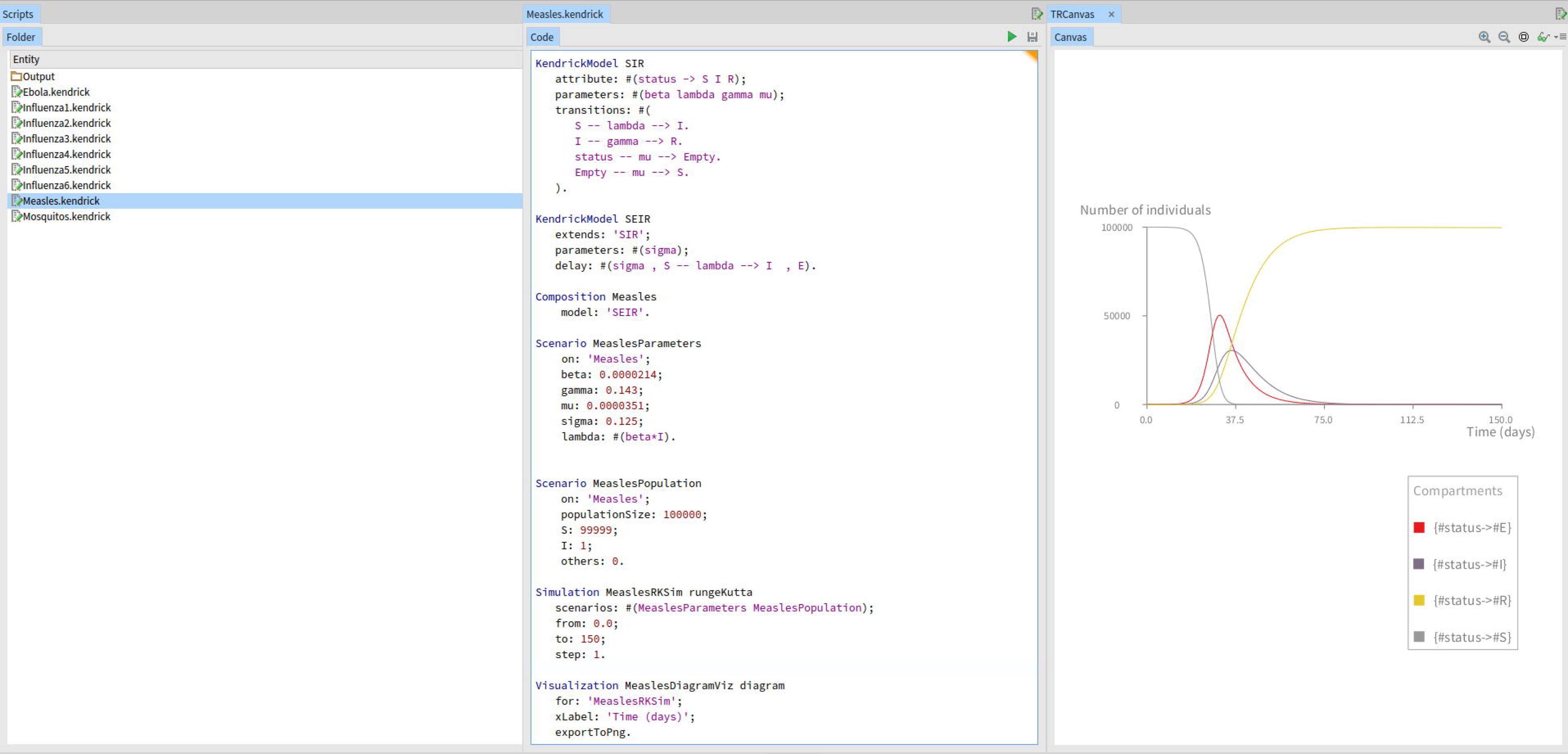
Kendrick UI. Running the Measles Script

The scripting version of the model is shown below:

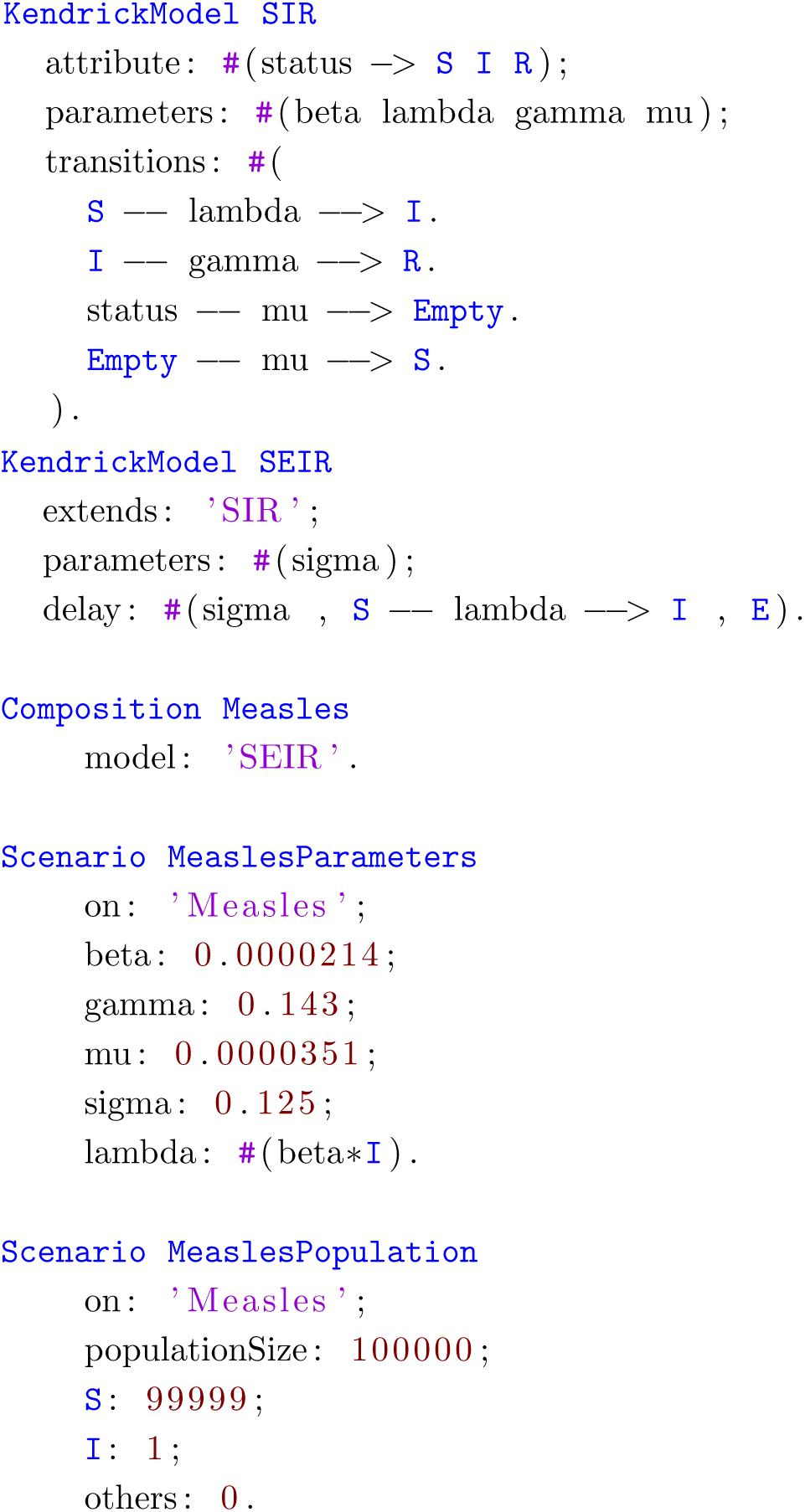

Where we can see lines 1 to 5 defining the SIR Epidemiological concern and the named transition parameters between its compartments. Then from lines 11 to 14 the SIR concern is extented to produce the SEIR compartmental description by introducing a delay parameter (sigma) and a compartment (E) between S and I. Subsequently, in lines 16 to 19 the Measles model is produced by using the SEIR concern while defining the formula for computing lambda. Complete models (such as Measles) are produced by the Composition entity of the DSL which as we saw on Table 1 can combine several different modelling concerns. Finally, from lines 21 to 33 two experiment scenarios are declared (one describing parameter values and the other population data) each of which can be used independently and combined with other initialization descriptions.

This version of the Measles model relies on the transition syntax of **Kendrick**, but alternative formalisms such as ODEs are also possible as seen below:

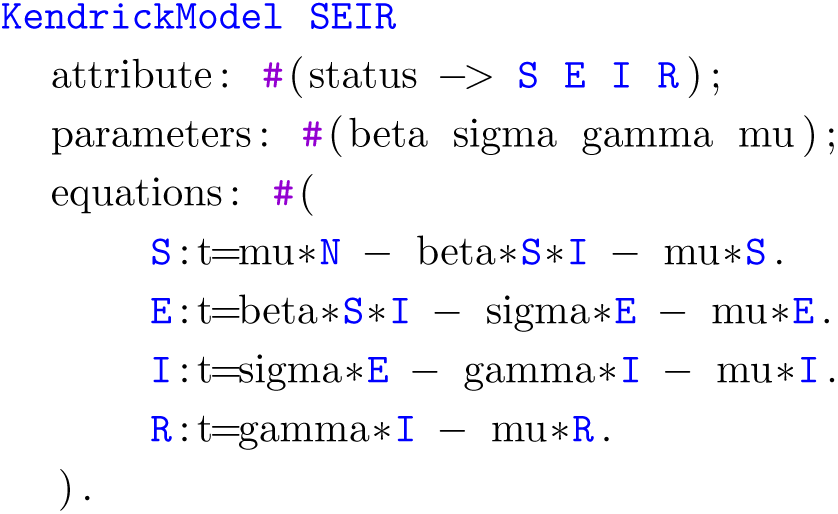

We should note here that frequently used entities such as SIR or SEIR shown above are part of the standard **Kendrick** library (available through our editor by navigating to Library/KendrickModel, seen in Figure 4).

**Figure 4.**
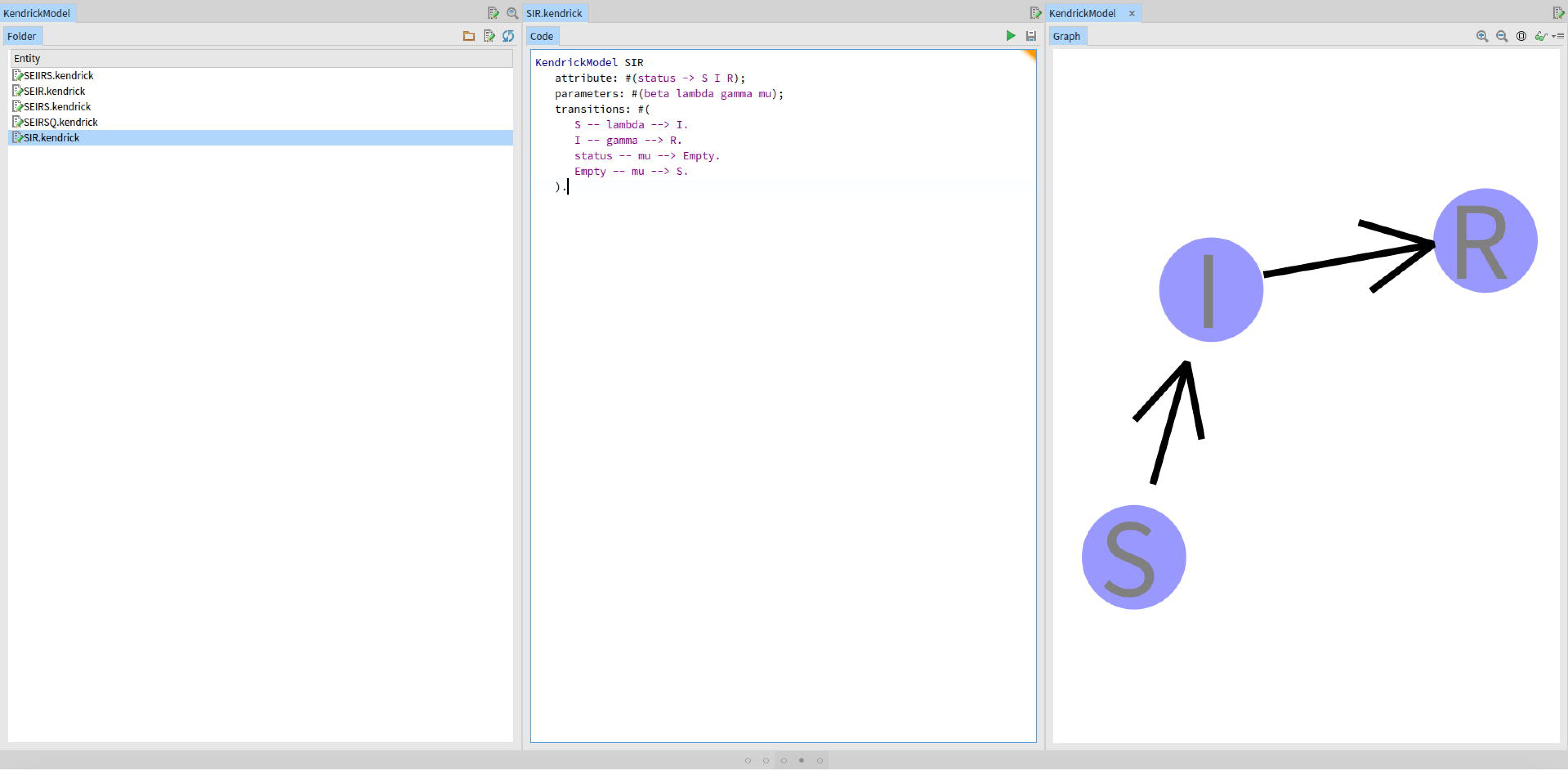
Kendrick UI. The Entity Library

The initial size of the population is 100,000 individuals and the parameters above, are taken directly from the related literature [18, 38]. Figure 5 shows the result of the deterministic simulation (using the RK4 solver) produced by the Simulation and Visualization entities shown below:

**Figure 5.**
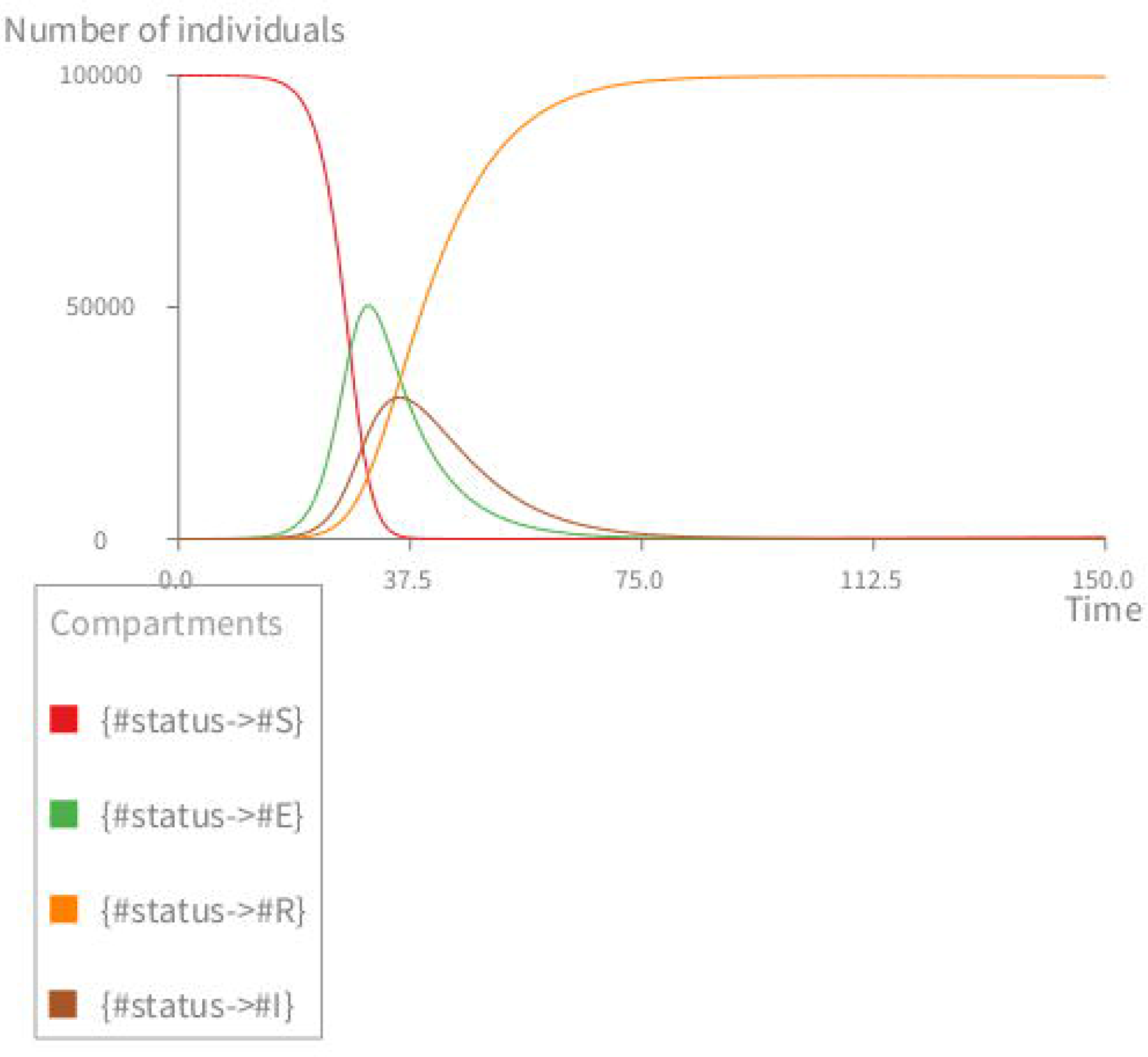
Deterministic simulation of the measles model. *S* = 99999, *E* = 0, *I* = 1, *R* = 0, *β* = 0:0000214, 1/*γ* = 7 days, 1/*σ* = 8 days, *ε* = 1/(78 * 365) in day^−1^, *N* = 100000. The graph shows the infectious deterministic dynamics of the measles model using the Kendrick language.

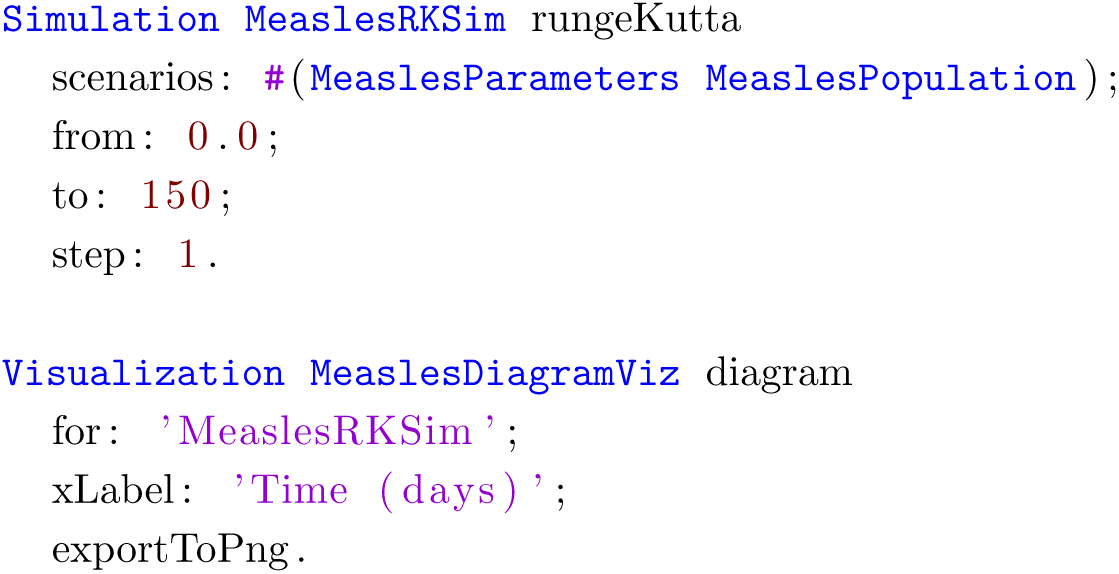

The Simulation entity (from lines 1 to 5) uses the two scenarios we defined previously, in addition of defining the specific algorithm to be used (rungeKutta) and the timing and stepping charecteristics. Finally, the Visualization entity specifies the desired output for a specific Simulation and by default will plot the dynamics of the Infected compartment over time.

### Case-study II: Mosquito-borne disease with three host species

As a second example we will study an SIR model with demography of a mosquito-borne disease with three host species. These species include a vector species (mosquito) and two potential host species species, designated as reservoir1 and reservoir2. The population is partitioned using two attributes: *status* and *species* leading to an arrangement of 3x3 compartments.

All species have the same six transitions: birth, deaths (for each one of the three status compartments), infection and recovery. Given a transition, the probability function is the same for everyone. Only 4 generic transitions need to be defined with **Kendrick**, which will then automatically generate the remaining transitions for each species of the model.

As previously the model is available through the standard **Kendrick** distribution by navigating through our editor either to Scripts/Mosquitos.kendrick (single file script, seen in Figure 6) or to Projects/Mosquito.

**Figure 6.**
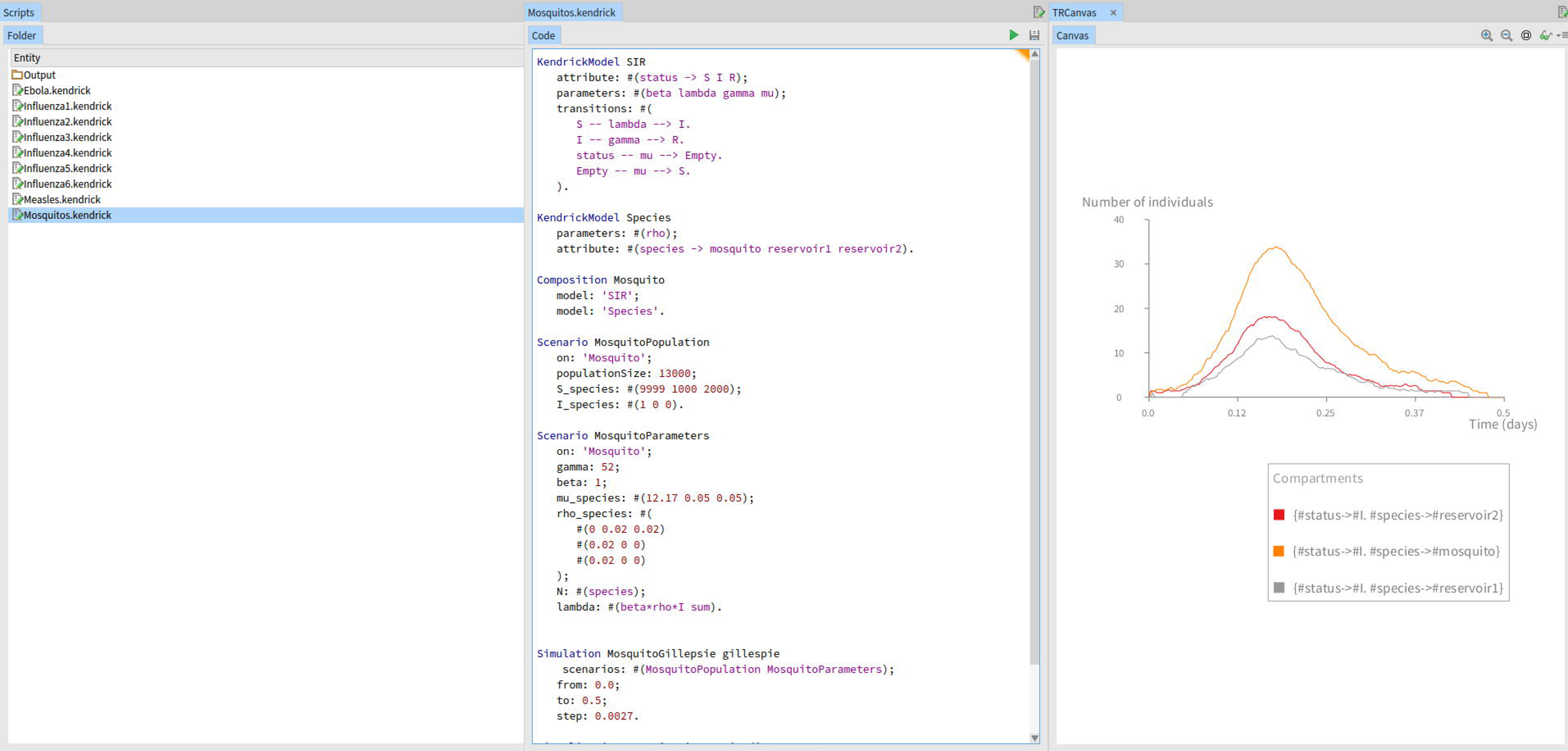
Kendrick UI. Running the Mosquitos Script

The scripting version of the model is shown below.

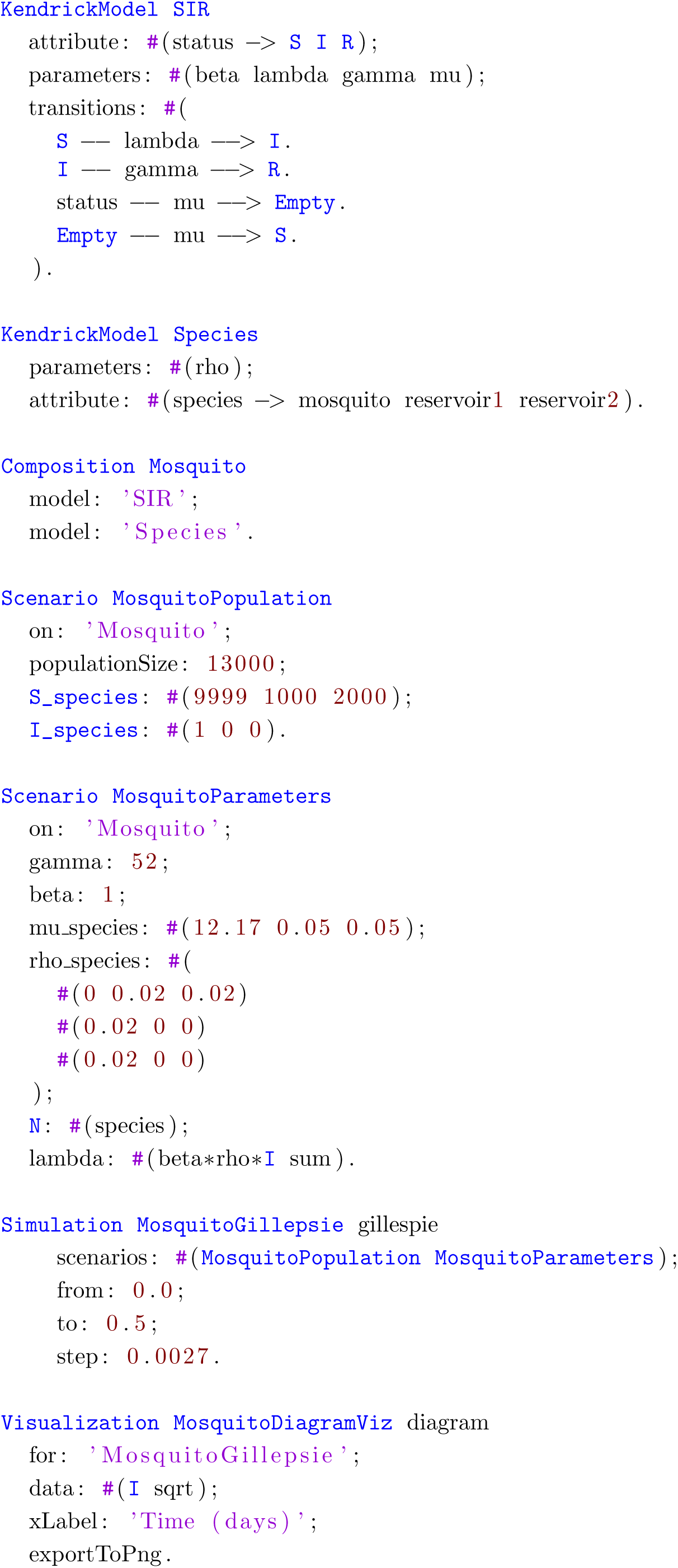

Lines 1 to 9 define the SIR example with demography (also available as a library model). Then on lines 11 to 13 the ecological concern is described, with the 3 species mentioned above, including the related rho parameter of our model. The composition of the epidemiological and ecological concerns takes place on lines 15 to 20, where the population N and the lambda parameters are declared. Two scenario entities follow (on lines 22 to 37) covering the population initialization for the different compartments, as well as the concrete parameter values for the rest of the model. The Simulation entity uses the two scenarios defined above to initialize the Gillespie algorithm for the given time-frame and stepping characteristics. Finally, the Visualization entity specifies the infectious stochastic dynamics to be plotted, over time (as seen on Figure 7).

**Figure 7.**
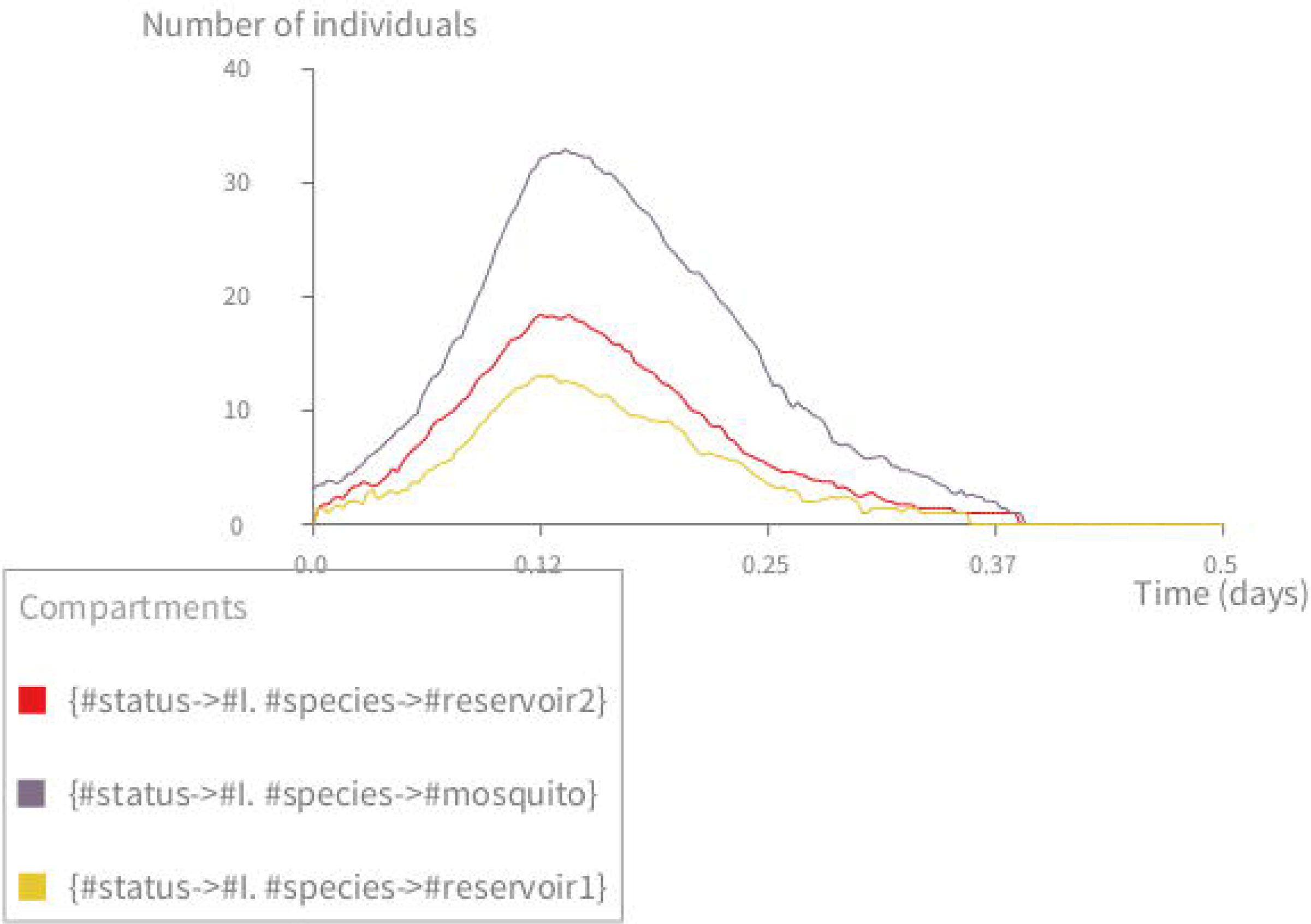
Stochastic simulation of the mosquito-borne disease. The multi-host model with three species: mosquito, reservoir1, reservoir2. *S*_1_ = 9999 (mosquito), *S*_2_ = 1000 (reservoir 1), *S*_3_ = 2000 (reservoir 2); *I*_1_ = 1, *I*_2,3_ = 0, *R*_1,2,3_ = 0, *N*_1_ = 10000, *N*_2_ = 1000, *N*_3_ = 2000; *σ* = 52, *µ*_1_ = 365/30, *µ*_2,3_ = 1/20, *β*_12_ = *β*_13_ = 0.02, *β*_*others*_ = 0.0.

### Validating the implementation of our language and simulation platform

To validate the core functionalities of **Kendrick** as a modelling and simulation platform, we first compared the output of deterministic **Kendrick** models with reference implementations of the same models on Scilab [39]. The simulation results (as those shown in Figure 5) suggest that for deterministic models **Kendrick** produces identical results to the reference implementations.

The dynamics of deterministic simulations have also been compared with those of Gillespie-based and agent-based simulations (Figure 8).

**Figure 8.**
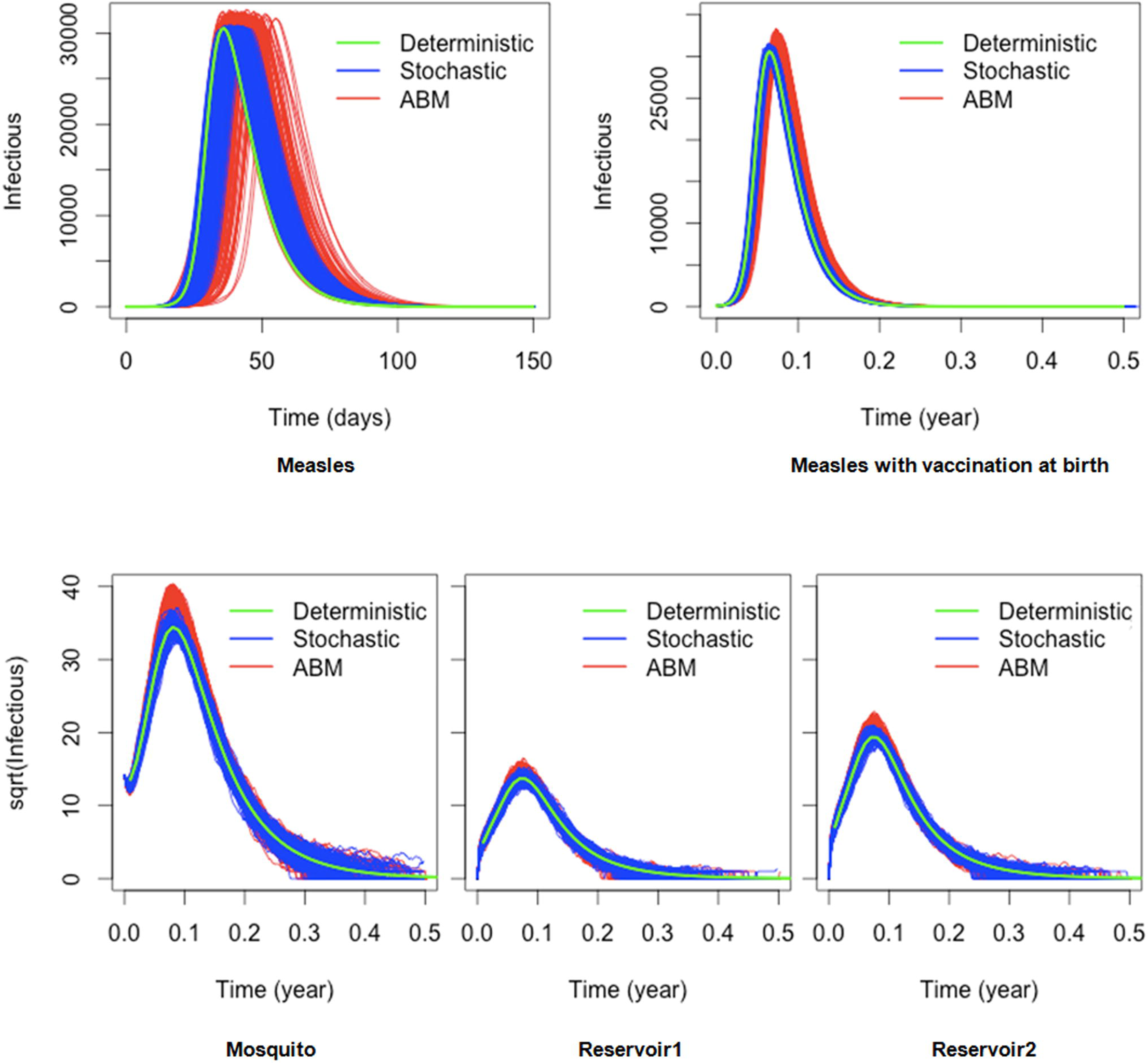
Comparison between the dynamics of deterministic, stochastic and agent-based model. We show the simulation results of two models in three formalisms: deterministic (green line), stochastic (blue lines) and agent-based (red lines). The first row shows the results of the measles model. The measles model is on the left hand side and the vaccination model is on the right hand side. The second row shows the results of the mosquito-borne model with three host species

The results for the measles model (taking into account scenarios with and without vaccination) can be seen in the upper part of Figure 8, with the lower part displaying the results of the mosquito-borne model for each individual host species. The deterministic dynamics can be superimposed on the stochastic and agent-based ones, validating our implementation of the stochastic and agent-based simulation logic.

Finally, we cross-examined agent-based and stochastic simulation outputs with the same configuration. Given that each agent-based and stochastic model is executed 200 times, we extracted from the simulation results key properties of the epidemiological dynamics, such as: epidemic peak, time at epidemic peak, and epidemic duration. A Kolmogorov-Smirnov statistical test on each pair of samples confirmed that there is no statistical difference between the agent-based simulations and the stochastic ones. These test results can be seen in Table 2, where we observe all *P − values* being greater than 0.05, validating that the resulting distributions are statistically indistinguishable. This in turn shows us the expected equivalence of the two algorithms, re-affirming the correctness of their implementation.

**Table 2.** P-values of Kolmogorov-Smirnov Test on two models over some disease global properties Model Global properties of diseases

| Model |  | Global properties of diseases |  |  |
| --- | --- | --- | --- | --- |
|  |  | Peak of epidemic | Time at peak of epidemic | Epidemic duration |
| Measles | Without Vaccination | 0.8762 | 0.2471 | 0.5246 |
|  | With Vaccination | 0.8928 | 0.2467 | 0.6262 |
| Mosquito-borne disease | Species 1 | 0.7920 | 0.2700 | 0.7112 |
|  | Species 2 | 0.6272 | 0.3927 | 0.8643 |
|  | Species 3 | 0.7920 | 0.1122 | 0.2202 |

## Discussion

Although DSLs have been used before in the context of bioinformatics [41, 40, 42], only a small number of them focused on epidemiological modelling [43, 44]. For example, *Ronald* [43] is a DSL for studying the interactions between malaria infections and drug treatments, but has unfortunately been discontinued. Schneider and al. [44] also proposed a DSL for epidemics, but their solution only supported agent-based models. Mathematical modelling languages (MMLs) such as Scilab [39], Modelica [45], Matlab [46] or JSim [47] do allow easier definition of mathematical models as sets of ODEs, but are too broad in scope to properly cover the domain-specific needs of epidemiology. On the other hand, individual libraries targeting epidemiological modelling: such as Epipy - a visualisation data tool for epidemiology written in Python [48], or GillespieSSA - an R package for generating stochastic simulation using Gillespie’s algorithms [49], cover only very specific sub-problems and epidemiological needs.

Closer to our approach are computational modelling tools for epidemiology such as FluTe [50], GLEAMviz [51], STEM [52] and FRED [53]. These solutions use dedicated approaches to model the transmission of infectious disease and provide a graphical user interface (GUI) to specify and visualise an epidemiological model. The main features of these tools are summarised in Table 3 and compared to **Kendrick**.

**Table 3.**
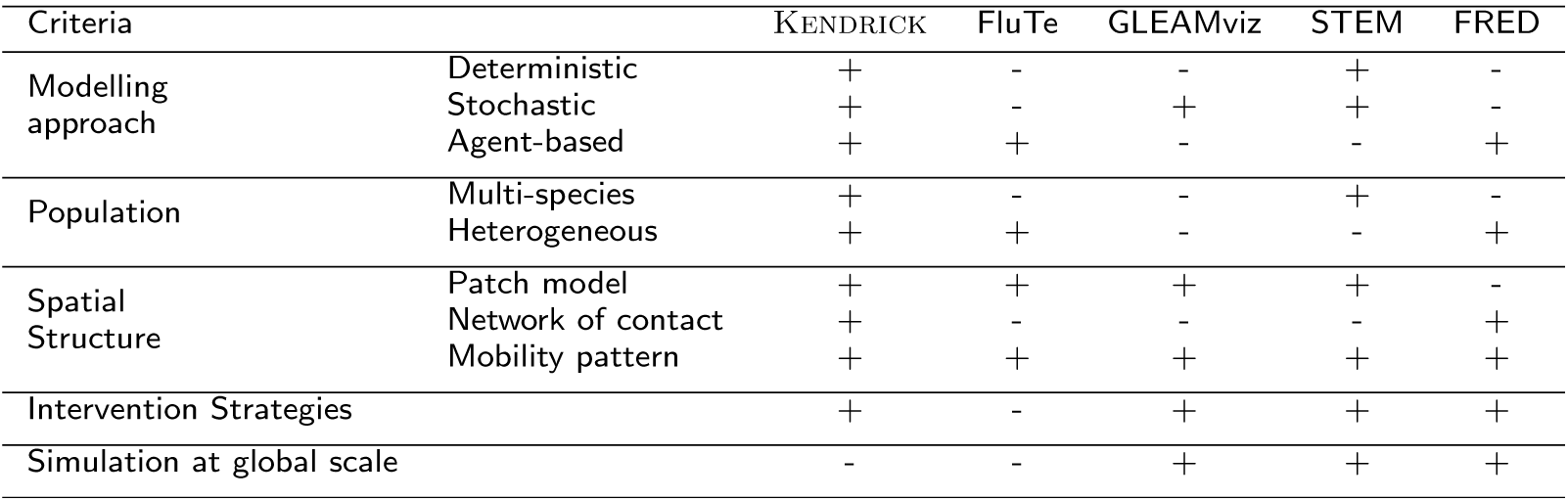
Modelling and Simulation Tools for epidemiology

Contrary to solutions that only focus on stochastic simulations (such as GLEAMviz [51] and STEM [52]) our platform can provide more detailed modelling thanks to the availability of agent-based simulation. Investigation of epidemics at a global scale as provided by GLEAMviz [51], STEM [52] or FRED [53] is also possible in **Kendrick**. A case study of meta-population model can be found in the additional file S1 provided with this publication. This particular case is also an example of spatial visualisation integrated within the platform.

Given the above comparison, **Kendrick** can be considered as a higher-level disciplined solution that aims to cover as many specificities of epidemiological modelling as possible. The fact that **Kendrick** chooses the right level of abstraction for each case, also helps modellers to focus on what is essential and avoid irrelevant or inconsistent definitions. This allows for easier cross-examination of different modelling paradigms (such as deterministic, demographic-stochastic and agent-based). From the modellers point of view, working directly with epidemiological concepts such as compartments, is more familiar than the underlying mathematical abstractions and algorithmic implementations.

In order to provide these abstractions in a transparent way for the user, our core model divides the population of **Kendrick** models in homogeneous classes, using mathematical equivalence relations (described in more details in [54]). By doing so we are able to capture different scenarios of infectious diseases such as multi-hosts, multi-strains[55, 21], heterogeneity, meta-population[2, 10] or network of contacts[1].

## Conclusion

In this paper, we have introduced a modelling language and simulation platform that allows specifying and simulating epidemiological models. The goal of our modelling approach is to allow experts to focus on their conceptual vocabulary through a dedicated DSL for epidemiology. We have highlighted the capacities of our platform through two classic examples of epidemiological models and we showed that the dynamics produced by our software are equivalent to well-established but harder to use, programming platforms.

By generalising the domain concepts used by experts, we have constructed a general language meta-model for epidemiology. This allowed us to express various epidemiological concerns such as spatial transmission, mobility, heterogeneous populations, evaluation of control strategies, visualisation, etc… in a uniform manner. We thus hope that our platform will be further adopted and extended to consider even more facets of epidemiology.

The **Kendrick** platform is available as an open source software under the MIT licence: http://ummisco.github.io/kendrick/.

## 1 Declarations

Availability of data and materials

Project name: KENDRICK

Project home page: http://ummisco.github.io/kendrick/

Operating System: multi-platform (Linux/Mac OS X/Windows)

Programming environment: Pharo 6.1: http://www.pharo.org/

Requirements: All the required tools for the installation of KENDRICK are described on the project home page.

License: MIT License

## Authors’ contributions

BTMA participated in study conception, carried out the implementation of the software, carried out the experimentation of the models, analysis of simulation results and drafted the manuscript. NP participated in study conception, carried out the DSL implementation and improved the manuscript. BR participated in study conception, to carry out the experimentation of models, results analysis and improved the manuscript. SS participated in study conception, carried out the implementation of the software and improved the manuscript. MZ participated in study conception and improved the manuscript. All authors read and approved the final manuscript.

## Acknowledgements

This work was supported by Alexandre Bergel (Chile University) and his team Object Profile (http://objectprofile.com/ObjectProfile.html) with the visualisation tool ROASSAL. We would like to acknowledge the financial support from Agence Nationale de la Recherche (ANR) PANIC project and European Smalltalk User Group (ESUG). We also gratefully acknowledge the financial support from Hanoi University of Science and Technology through the “Spatial Modelling Language for Epidemiology” project - grant number T2017-TT-001.

## Additional Files

Additional file 1 (s1) — The meta-population model with immigration infection specified in Kendrick language

## Footnotes

[1] https://github.com/UMMISCO/kendrick

[2] http://smalltalkhub.com/#!/~UMMISCO/Kendrick

[3] https://travis-ci.org/UMMISCO/kendrick

[4] http://ummisco.github.io/kendrick/

[5] https://github.com/UMMISCO/kendrick/wiki

[6] https://github.com/UMMISCO/kendrick

[7] http://designinginterfaces.com/firstedition/index.php?page=Cascading_Lists

